# DeepQSM - Using Deep Learning to Solve the Dipole Inversion for MRI Susceptibility Mapping

**DOI:** 10.1101/278036

**Authors:** Kasper Gade Bøtker Rasmussen, Mads Kristensen, Rasmus Guldhammer Blendal, Lasse Riis Østergaard, Maciej Plocharski, Kieran O’Brien, Christian Langkammer, Andrew Janke, Markus Barth, Steffen Bollmann

## Abstract

Quantitative susceptibility mapping (QSM) aims to extract the magnetic susceptibility of tissue from magnetic resonance imaging (MRI) phase measurements. The mapping of magnetic susceptibility *in vivo* has gained broad interest in several fields of science and medicine because it yields relevant information on biological tissue properties, predominantly myelin, iron and calcium. Thereby, QSM can also reveal pathological changes of these key components in devastating diseases such as Parkinson’s disease, Multiple Sclerosis, or hepatic iron overload. As QSM requires the solution of an ill-posed field-to-source-inversion, current techniques utilize manual optimization of regularization parameters to balance between smoothing, artifacts and quantification accuracy. We trained a fully convolutional deep neural network - DeepQSM - to invert the magnetic dipole kernel convolution. This network is capable of solving the ill-posed field-to-source inversion on real-world *in vivo* MRI phase data without the need for manual parameter tuning, which proves that this network has generalized the underlying mathematical principle of the dipole inversion. We demonstrate that DeepQSM’s susceptibility maps enable identification of deep brain substructures that are not visible in MRI phase data and provide information on their respective magnetic tissue properties. We illustrate DeepQSM’s clinical relevance in a patient with multiple sclerosis showing its sensitivity to white matter lesions. In summary, DeepQSM can be used to determine the composition of myelin sheets of nerve fibers in the brain, and to assess quantitative information on iron homeostasis and its dysregulation, and will subsequently contribute to a better understanding of these biological processes in health and disease.

## Introduction

Quantitative susceptibility mapping (QSM) is an increasingly utilized post-processing technique that extracts magnetic susceptibility from the phase of magnetic resonance imaging (MRI) gradient echo signal (1, 2). Magnetic susceptibility describes the degree of magnetization of a material placed in an external magnetic field and thereby delivers unique, noninvasive insights into tissue composition and microstructure (3, 4). In particular, QSM provides information about myelin and white matter composition (5, 6), iron metabolism (7– 12), and copper accumulation (13). The measurement of iron stores has been used to study normal aging (7, 14), Huntington’s Disease (8), Multiple Sclerosis (15–17), Alzheimer’s Disease (18) and Parkinson’s Disease (19, 20). Furthermore, QSM visualizes micro-bleeds (21) and differentiates them from microcalcifications (22) due to differing magnetic susceptibilities of calcium and iron.

In order to obtain a quantitative susceptibility map, an image is acquired using an MRI sequence where the signal phase is sensitive to local magnetic field changes, such as a gradient-recalled echo sequence (23–25). This raw signal phase is unwrapped and magnetic field contributions from outside the object of interest, the so-called background field, are removed. Finally, the inverse problem relating the measured field perturbation to the underlying magnetic susceptibility distribution is solved (1, 2). This critical inversion step is ill-posed, because multiple susceptibility distributions could result in the same field perturbation. The ill-posed nature can be overcome either by additional measurements in different orientations or by numerical stabilization strategies.

One method utilizing the acquisition of different object orientations with respect to the static magnetic field is known as ‘Calculation of susceptibility through multiple orientation sampling’ (COSMOS (26, 27)). COSMOS requires at least three different orientations to make the field-to-susceptibility problem over-determined and solves the inverse problem analytically. Although COSMOS generates almost artifact-free susceptibility maps, and is therefore seen as gold standard, it assumes isotropic magnetic susceptibility and contains little information about anisotropic tissues (28, 29). Therefore, methods such as susceptibility tensor imaging (STI) (30) or the Generalized Lorentzian Tensor Approach (GLTA) have been developed that extend the magnetic susceptibility scalar to a tensor. Common to all multi-orientation methods is their clinical in-feasibility due to patient discomfort and scan time requirements (1, 31). To overcome these practical limitations, a variety of methods have been developed that compute magnetic susceptibility from single orientation data by employing numerical stabilization techniques.

Numerical strategies can be subdivided into two groups (2): inverse filtering and iterative methods. Inverse filtering formulates the problem in Fourier domain where dividing the pre-processed phase data by the unit dipole response yields the magnetic susceptibility. However, near-zero values in the unit dipole response result in an amplification of noise and errors - a manifestation of the ill-posed problem. The unit dipole response is therefore modified and small values are replaced by a fixed threshold. This method is known as truncated k-space division (TKD (32, 33)). Due to the now inaccurate physical model, TKD parameters need to be carefully chosen to yield a trade-off between regularization and artifacts and TKD results need to be corrected for underestimating magnetic susceptibility (2).

It is also possible to solve the inverse problem iteratively in the spatial domain (2). Such iterative numerical solvers require the description of the forward solution (the multiplication of the dipole kernel with the susceptibility distribution in Fourier space) and they minimize the difference between this predicted data and the actual data in a least-squares sense. One example to solve this equation system is the LSQR algorithm (34, 35). Most QSM inversion algorithms developed in the last years are extensions to this basic principle but differ in the way how regularization techniques incorporate prior information about the susceptibility distribution. Common to all techniques is that the regularization terms have to be carefully optimized to yield a trade-off between data and priors (2, 36). Morphology enabled dipole inversion (MEDI) is a spatially regularized inversion that utilizes edge information from magnitude images to regularize the problem (37, 38). Other methods include total generalized variation (TGV (39)), or single-step QSM (SS-QSM (40)). Common to all iterative methods is that the forward solution has to be evaluated at least once in every iteration step. This results in two consequences: 1) Iterative methods are considerably slower than inverse filtering methods and 2) the forward solution has to be simple enough to be computed in every iteration.

Recently, deep neural networks have emerged as an alternative to iterative methods for solving inverse problems (41– 44) and have shown impressive results in applications such as denoising (45), deconvolution (46), image reconstruction (47–49) and super-resolution (50–53). The use of neural networks for the solution of inverse problems is motivated by the fact that neural networks are capable of approximating any continuous function under the assumption that the network has enough free parameters (42, 44, 54). Researchers have also investigated the link between iterative methods and deep networks and found that neural networks act as a fast approximated sparse coder (41, 55). A sparse coder learns the most important features from the input data and can reconstruct the input using a combination of basis vectors (55). It has been shown (41) that the reconstructed images from the neural network preserved complex textures better than the state-of-the art iterative method, because the neural network learns an efficient regularization from the data, whereas iterative methods require explicit regularization. An important practical advantage of neural networks over iterative methods is that the trained models can produce an output extremely fast as they only involve a single step of matrix multiplications to produce the feed forward output (42).

Based on the recent advances in the solution of inverse problems using deep learning, we propose a deep convolutional network that delivers fast and artifact-free solutions to the ill-posed field-to-source problem. We test the generalizability in four separate experiments with increasing complexity: The first experiment tested the performance on a data set very similar to the training data. The second experiment investigated the network’s performance on a more realistic brain data set, containing structures that the network has never seen during training. The third experiment goes one step further and tests if the inversion works with real-world *in vivo* phase data of a healthy participant. Finally we test the performance of our algorithm on a clinical dataset from a patient with multiple sclerosis. We show that DeepQSM is capable of utilizing real-world single-orientation phase data without the need for explicit regularization terms and manual parameter tweaking. The solutions are fast, robust and show a high level of detail (see Figure 1).

**Fig. 1.**
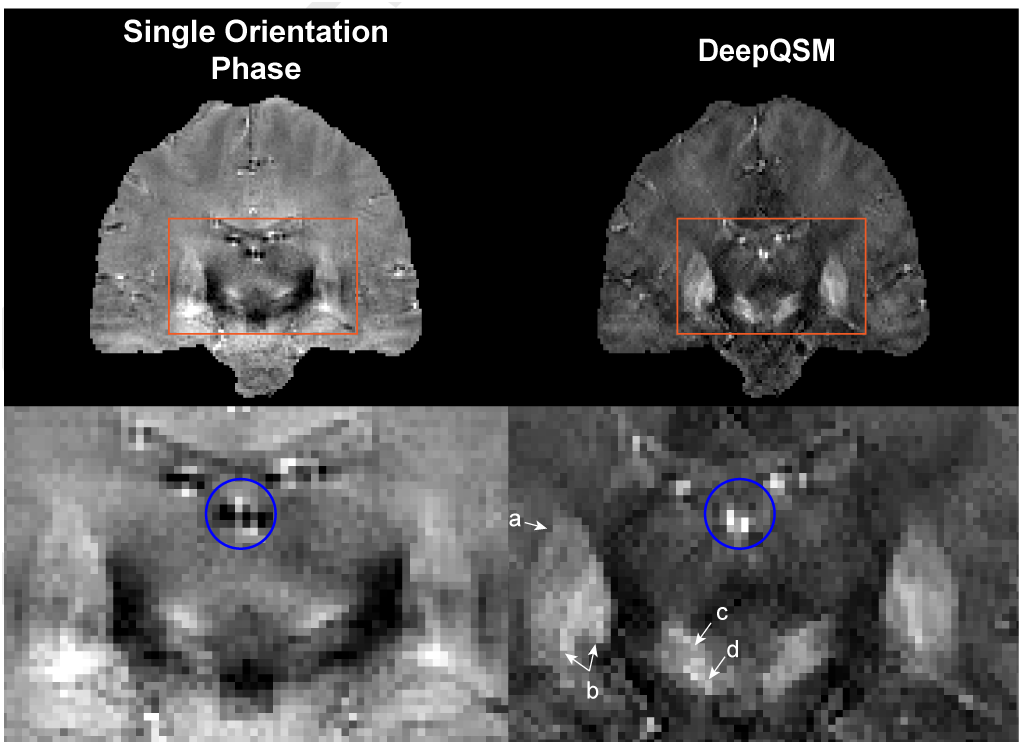
DeepQSM’s susceptibility maps enable identification of deep brain substructures. In this particular coronal slice there are indications of the following substructures: **a)** *putamen*, **b)** *globus pallidus*, **c)** *subthalamic nucleus*, **d)** *substantia nigra*. The left column shows the same slice of the single orientation phase image used for susceptibility reconstruction. The blue circle highlights a dipole artifact around a vein that is removed in the QSM reconstruction.

## Methods

### Network Architecture

The fully convolutional neural network (DeepQSM) capable of processing 3-dimensional inputs is based on a modified version of an established architecture (U-Net) (56). Due to memory constraints on the Graphic Processing Units (GPUs) used for training we reduced the amount of feature-maps compared to the original U-Net, see Figure 2 for the architecture of DeepQSM. The fully convolutional nature of the chosen architecture allows an input image of any size with dimensions divisible by 16 and the dimensions of the output equal the dimensions of the input. The architecture can be divided into a contracting and an expanding part. The goal of the contracting part is to capture the context of the image, while the goal of the expanding part is to increase the resolution (41). The contracting part of DeepQSM consists of three-dimensional convolution layers with filters of size 3×3×3, a stride length of 1×1×1 and rectified linear units (ReLU). Furthermore, pooling layers are added which both increase the receptive field and make the network translation invariant. The expanding part of the neural network consists of two types of convolutional layers: transposed convolutional layers and convolutional layers. The transposed convolutional layers consist of filters of size 2×2×2, a stride length of 2×2×2 and ReLUs. The convolutional layers in the expanding part are identical to the convolutional layers in the contracting part. The network has skip connections between the contracting and expanding part of the network. The skip connections make up for the spatial information lost during downsampling by combining high resolution information with low resolution information. Furthermore, it reduces the gradient vanishing problem and increases the performance of the network (41).

**Fig. 2.**
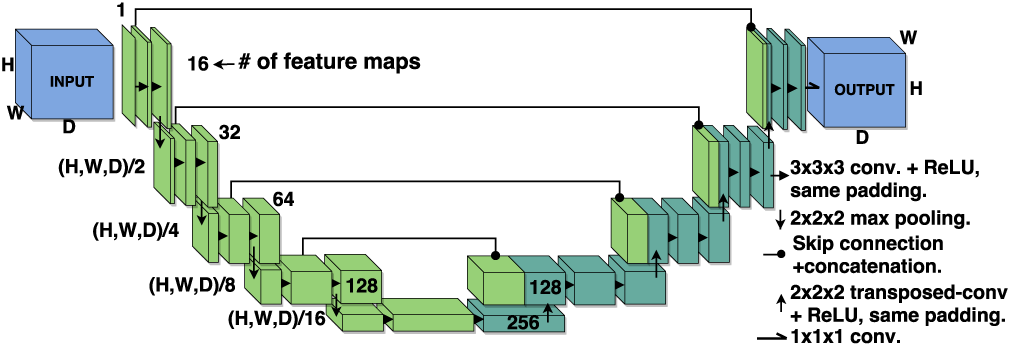
The DeepQSM architecture consists of a contracting and expanding part. The contracting part is made up of a series of convolutions with a ReLU activation function followed by a pooling layer. The expanding part consists of transposed convolutions to undo the spatial reduction caused by the pooling operations and convolutions with ReLUs similar to those of the contracting part. Convolutions used a stride of 1×1×1, transposed convolutions a stride of 2×2×2, pooling a stride of 2×2×2. The input given to the image must have spatial dimensions that yield a positive integer when divided by 16. DeepQSM will output a volume with identical dimensions to the input.

### Dipole Kernel

The dipole kernel accounts for the fact that susceptibility is a non local property. The kernel commonly used in QSM (1, 2, 40, 57) is built on two assumptions. The first assumption is that the effect of the local environment on susceptibility can be divided into near field and far field based on the Lorentz sphere (58). Under the assumptions that the magnetic susceptibility is a bulk property and that magnetic moments in the near field are randomly distributed, the contribution of the near field can be neglected (58). Equation 1 is the dipole kernel in the Fourier domain, where *k*_*x*_,*k*_*y*_ and *k*_*z*_ are k-space vectors in the respective directions. This dipole description does not include terms for modeling anisotropy:

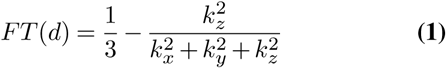

### Training Procedure

DeepQSM was trained on 70400 48×48×48 synthetic examples in 20500 steps with 32 examples per step resulting in a total training time of 15 hours. The synthetic 3D images were simulated and contained basic geometric shapes such as cubes, rectangles and spheres and served as ground truth/label data. Input data was generated from the label data by convolving it with the dipole kernel (see eq. 1) to create well-posed forward solutions (see Figure 3). To optimize the weights of DeepQSM during training the ‘ADAM’ optimizer (59) was used, and had the following configurations: initial learning rate = 0.001, *β*1 = 0.9, *β*2 = 0.99. Mean squared error between the reconstruction from DeepQSM and the label data served as the cost function for the optimizer.

To avoid overfitting the weights during training and thereby losing generalisability, the regularization technique ‘dropout’ was used (60) to randomly turn off neurons during training with a drop out rate of 10%. DeepQSM was implemented using Python 3.6.1 and Tensorflow v1.3.1. Training was performed on the National Computational Infrastructure cluster ‘*Raijin*’ using NVIDIA Tesla K80 GPUs.

### Test

Four experiments were performed to evaluate DeepQSM’s ability to solve the ill-posed field-to-source inversion. Each experiment tests DeepQSM’s performance and generalisability by gradually increasing the complexity of the data used for each experiment. Figure 3 shows how data was processed before a DeepQSM reconstruction. For the first experiment synthetic data was used, while for the second and third experiment a single orientation background field corrected tissue phase image and STI susceptibility map published by (31) for *the 2016 QSM reconstruction challenge* were used. The data set serves as a common reference for current and future algorithms and was acquired *in vivo* from a healthy 30 year old female, using a 3D gradient-echo at 3T with 1.06mm isotropic resolution, an echo time of 25ms and a repetition time of 35ms (31). The phase data was obtained from a single orientation while the ground truth susceptibility map was computed using STI *χ*33 (30) from 12 orientations. The fourth experiment involved clinical data from a female patient with multiple sclerosis (30 years of age, Clinically Isolated Syndrome (CIS), Kurtzke Expanded Disability Status Scale (EDSS) = 2.0). The 3D gradient-echo data were acquired on a 3T with 1×1×2mm resolution and 6 echos with echo times of 4.92ms to 29.2ms.

**Fig. 3.**
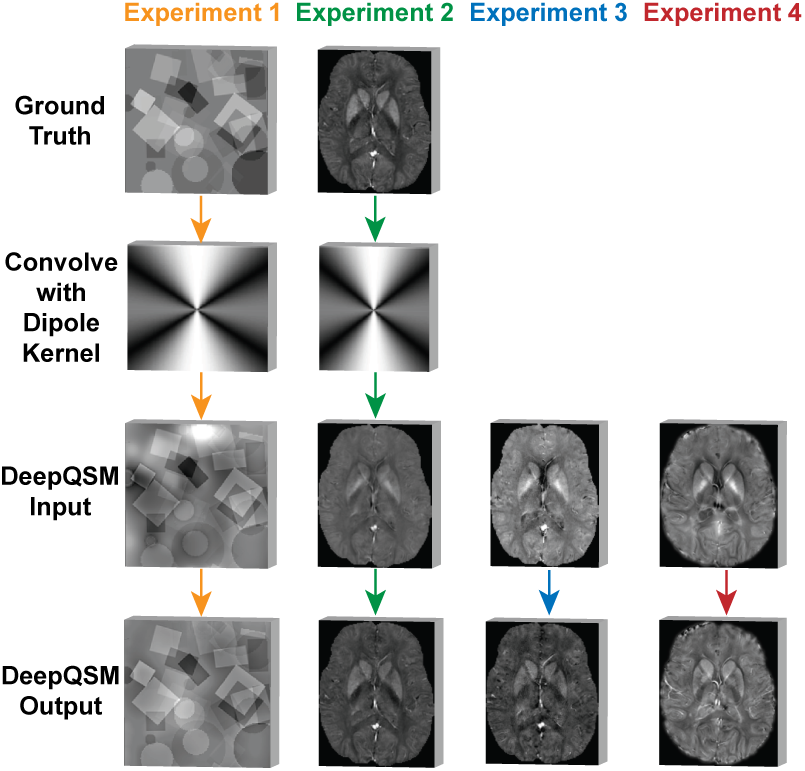
Illustration of the data-flow for experiment 1 (orange arrows), 2 (green arrows), 3 (blue arrows) and 4 (red arrows). Experiment 1 and 2 gives the well posed forward solution calculated by convolving the ground truth with the dipole kernel as an input to DeepQSM. Experiment 3 and 4 use a background field corrected single orientation 3D GRE MRI phase as input for DeepQSM.

***Experiment 1.*** The first experiment aimed to test DeepQSM’s performance on synthetic data similar to the training data. The goal was to see if DeepQSM would produce sensible outputs on new synthetic data that simulated exactly the same way as data used for training, then convolved with the dipole kernel to generate a well-posed forward solution. Next DeepQSM was set to invert the applied dipole kernel to reconstruct the simulated susceptibility map.

***Experiment 2.*** In the second experiment the goal was to solve the ill-posed problem on a dataset dissimilar from the training data. DeepQSM had never been introduced to images of brains during training and therefore this experiment would test if DeepQSM generalized the underlying physical principle of the QSM dipole inversion. The forward solution was generated by using a brain *χ*33 solution as the ground truth and convolving it with the dipole kernel.

***Experiment 3.*** The third experiment aimed to test if DeepQSM can reconstruct a susceptibility map from realistic single orientation phase data and compare the reconstruction to the thresholded k-space division (TKD)(33), a widely utilized, fast, QSM reconstruction technique. TKD overcomes the ill-posed problem by setting values close to zero to a manually chosen threshold which depends on the noise and signal characteristics of the dataset. The threshold used for this data was 0.19 as it yielded the best-trade off between streaking artifacts and image quality.

***Experiment 4.*** The fourth experiment tests if DeepQSM can reconstruct a susceptibility map from clinical single orientation phase data of a patient with multiple sclerosis. The quantitative susceptibility maps are compared with a standard Magnetization Prepared Rapid Acquisition GRE (MPRAGE) and a fluid-attenuated inversion recovery (FLAIR) sequence.

## Results

We have trained a deep convolutional neuronal network on forward and inverse examples generated from synthetic data in 15 hours. The following experiments demonstrate the performance of the network trained on this synthetic data and applied to a variety of datasets with increasing complexity from experiment 1 to 4. For an illustration of the pre-processing and calculation of the results, see Figure 3. All DeepQSM predictions were computed in approximately 10 seconds on a standard workstation.

### Experiment 1: Synthetic Data

To verify that the network can solve an inverse problem similar to the training data, we applied the trained network to a synthetic dataset generated with the same rules as the training data. Figure 4 illustrates the prediction performance of DeepQSM on synthetic data and compares it to the simulated ground truth. This figure illustrates DeepQSM’s capability of reverting the effects of the dipole on synthetic data without suffering from streaking artifacts or introducing smoothing.

**Fig. 4.**
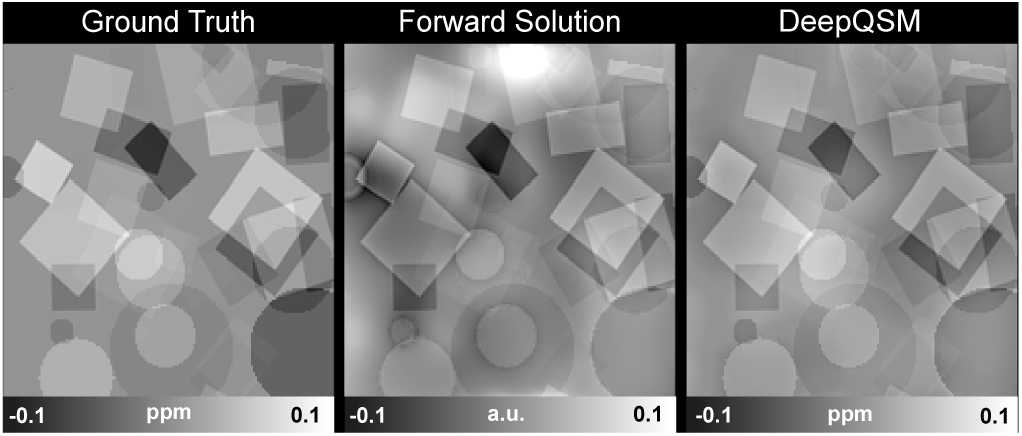
One representative slice from a 3D volume of synthetic data. Left: Ground truth, Middle: Forward solution obtained after convolving the QSM dipole kernel with the ground truth, Right: DeepQSM reconstruction of the forward solution.

### Experiment 2: Forward Simulation

The second experiment aims to test if the network can predict on structures dissimilar from data during the training-phase consisting of simple shapes. In this experiment we used a human brain QSM reconstruction from multiple orientations as the ground truth. This brain was then convolved with the dipole kernel to generate the input for DeepQSM (well-posed forward solution). Figure 5 shows DeepQSM’s ability to revert the ill-posed dipole kernel operation on a realistic brain dataset without introducing any artifacts.

**Fig. 5.**
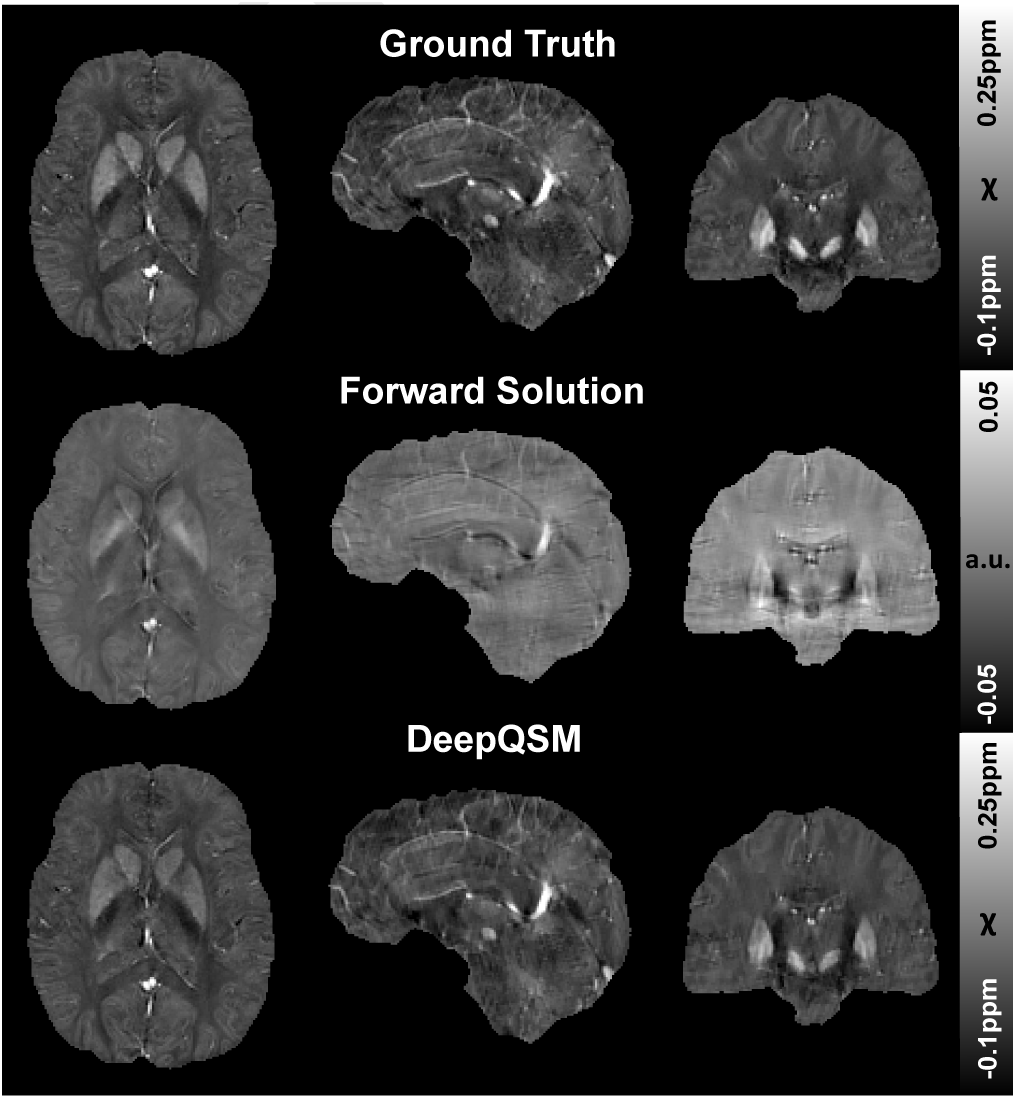
Axial, sagittal and coronal middle slice from a 3D volume. First row shows the ground truth, second row shows the result of convolving the ground truth with the dipole kernel to generate the input data for DeepQSM. The third row shows the resulting reconstruction by DeepQSM.

### Experiment 3: Single Orientation 3D GRE MRI Phase Data

The third experiment tests if DeepQSM can predict susceptibility maps based on measured single orientation *in vivo* phase data and compares the results to the established inversion technique TKD with a manually chosen threshold of 0.19 yielding the best trade-off between quantification accuracy and artifacts. The result can be seen in Figure 6 that shows how DeepQSM is able to solve the inverse problem on realistic phase data from a single orientation without introducing smoothing or requiring the explicit choice of regularization parameters.

**Fig. 6.**
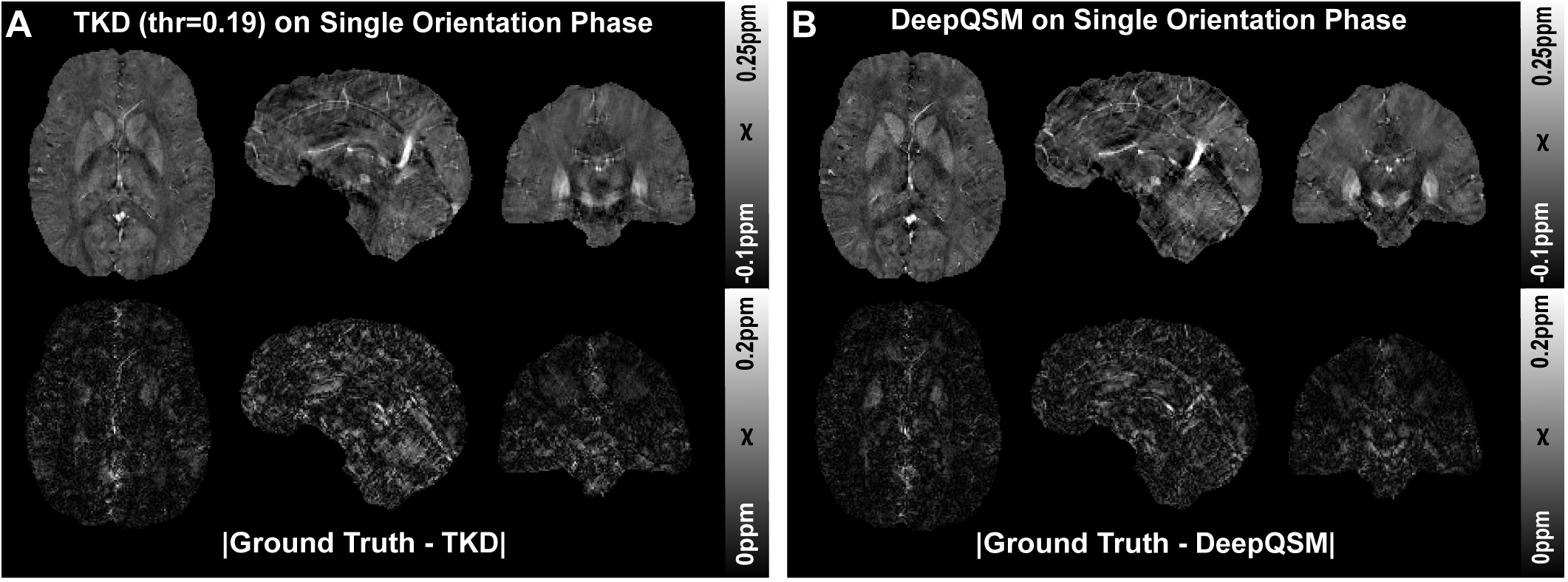
Comparison of **A**) thresholded k-space division (TKD) with a threshold value of 0.19. and **B)** DeepQSM. The first row shows an axial, sagittal and coronal slice of the reconstructed 3D susceptibility maps. The reconstructions have been performed on single orientation background field-removed phase data. The second row shows the absolute difference maps between the reconstructed image and the ground truth from the 2016 QSM reconstruction challenge.

A zoomed version of the coronal orientation is shown in Figure 1, to illustrate how substructures of interest such as *putamen, globus pallidus, subthalamic nucleus*, and *substantia nigra* are present in the susceptibility map resulting from DeepQSM.

### Experiment 4: Clinical 3D GRE MRI Phase Data

A clinical dataset is shown in Figure 7 to illustrate that DeepQSM can process real-world clinical data and show the sensitivity to multiple sclerosis lesions. This example also illustrates how little smoothing DeepQSM introduces with the consequence that small blood vessels are visible in the DeepQSM reconstruction.

**Fig. 7.**
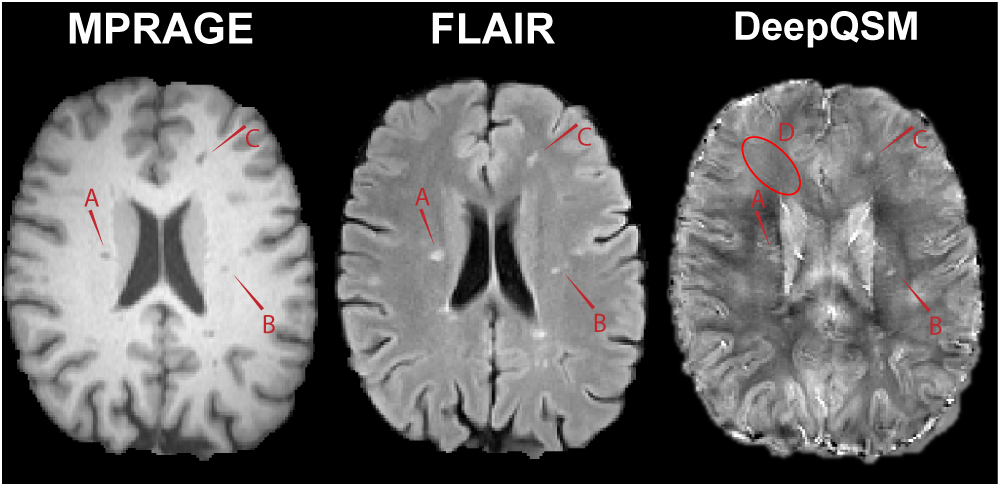
A clinical example showing data from a patient with multiple sclerosis. The image shows three modalities, MPRAGE (left), FLAIR (middle), and DeepQSM (right). The red arrows highlight lesions (A, B, C) and it can also be observed that DeepQSM allows distinguishing small blood vessels (D).

## Discussion

In this proof-of-concept study we demonstrated that the challenging inverse problem underlying QSM can be solved by using a fully convolutional neural network. DeepQSM yielded a low level of streaking artifacts and preserved small anatomical structures. We achieved this by computing the well-posed QSM forward solution of synthetic data consisting of simple shapes and we trained the network on these inverse and forward solution pairs. The first experiment tested the performance on a data set very similar to the training data. The network was able to invert this problem, but it could have done this by overfitting the training data due to its simple nature. Therefore, the second experiment investigated the network’s performance on a more realistic data set, containing structures that the network has never seen during training. We used the STI *χ*33 ground truth from the *2016 QSM reconstruction challenge* (31) and computed the well-posed QSM forward solution. Then we used the network trained on simple shapes to invert the field-to-source problem for a data set with realistic anatomical structures. This showed that the network has learned the general concept of the dipole inversion and not just the shapes presented during training. The third experiment goes one step further and tests if the inversion works with real-world *in vivo* phase data, containing measurement noise and error propagation through previous steps of the QSM pipeline. For this we used a single orientation scan from the *2016 QSM reconstruction challenge* (31) and demonstrate that the network can invert this problem while preserving fine spatial structures. Finally we show that deep-QSM can also invert clinical data from a patient with multiple sclerosis.

We compared the susceptibility reconstructions from DeepQSM with a widely utilized reconstruction method TKD and showed that DeepQSM is capable of delivering robust results. One of the major benefits of DeepQSM compared to TKD (and other existing algorithms) is that DeepQSM does not utilize explicit regularization parameters to yield a balance between smoothing, artifacts and quantification accuracy and additionally works as an end-to-end algorithm capable of computing susceptibility maps within seconds.

Although we used a very simplistic training data set we achieve high quality QSM reconstructions in the brain. This shows that our network has learned to approximate the underlying physics of dipole inversion and can potentially use training data sets with anatomical priors to help condition the ill-posed nature of the problem further. This could be achieved by utilizing a minimum deformation model (61, 62) of the human brain anatomy to generate high quality training data. The more similar the training data is to the real brain anatomy, the more prior knowledge the network will be able to utilize, similar to QSM algorithms exploiting morphological features (37).

We have demonstrated the potential of our newly presented approach by using brain scans. However, our method is not only applicable to solving the QSM inverse problem in the brain but can be extended to other regions in the body, such as the liver or the heart. For this, a basic network could be trained on simple shapes, as we did in this work, and then use this basic network in a second transfer learning phase, where the anatomical priors are learned from the organ of interest. It has been shown, that a fine-tuning of pre-trained convolutional neuronal networks outperforms networks trained from scratch (63). Fine-tuning a pre-trained network requires a smaller amount of training data compared to training a network from scratch (42). This is especially useful in the field of QSM where high quality training data is costly and difficult to obtain.

Currently, we have used background field corrected data to compute a QSM solution. However, it is also possible to incorporate realistic simulations of background fields originating at tissue boundaries into the training step. This would then allow the background field removal together with the field-to-source inversion in a single step, similar to state-ofthe-art iterative QSM algorithms (39).

In our current implementation we utilized a simple dipole model with the assumption that magnetic susceptibility is a scalar quantity. An advantage of our proposed approach is that it can potentially utilize any forward model and as such could incorporate additional model terms accounting for anisotropy of magnetic susceptibility and structural tissue anisotropy. Currently the inverse problem posed by these more sophisticated models, such as the Generalized Lorentzian Tensor Approach (GLTA) (58), cannot be solved with existing procedures. However, our deep learning approach could potentially incorporate additional measurements to help condition these inverse problems.

This new class of QSM algorithms has a wide range of clinical applications. DeepQSM could for example be combined with a fast imaging sequence based on echo planar imaging, such as the recently proposed planes-on-a-paddlewheel sequence (64). This would allow the routine acquisition and reconstruction of QSM data in under 1 minute, compared to current techniques requiring at least 5 minutes. This increases patient comfort and helps with motion artifacts in clinical populations. This could enable the standard clinical use of QSM in assessing microbleeds (21, 65), the diagnosis of Alzheimer’s (18, 66, 67), Parkinson’s (20, 68–72), and Huntington Disease (8), Multiple Sclerosis (17, 73, 74), Amyotrophic Lateral Sclerosis (75), Wilson Disease (13) or more general in diseases with a dysregulation of iron metabolism (12, 76, 77). QSM could also help in localizing targets, such as the Subthalamic Nucleus, for Deep Brain Stimulation Surgery (78).

## Conclusions

In summary, we have described the foundations for a new class of QSM inversion algorithms that allow the solution of the QSM inverse problem without requiring explicit regularization parameters and manual parameter tweaking. This has the potential to create algorithms that can reconstruct QSM from clinical single-orientation phase data in a fraction of the time that is currently required for standard iterative procedures. Our approach delivers artifact-free susceptibility maps and the presented algorithm can be extended to include more realistic forward models that could allow the modeling of anisotropic components in QSM.

## ACKNOWLEDGEMENTS

The authors acknowledge the facilities of the National Imaging Facility (NIF) at the Centre for Advanced Imaging, University of Queensland. The three first authors acknowledge funding from the following private organisations: Aalborg University Internationalisation foundation, Otto Mønsted foundation, Knud Højgaard founda-tion, Danish Tennis Foundation, Nordea foundation, Julie Damms study-foundation and Oticon foundation. CL acknowledges funding from the Austrian Science Fund (FWF grants KLI523 and P30134). SB acknowledges funding from UQ Postdoctoral Research Fellowship grant and support via an NVIDIA hardware grant. MB acknowledges funding from Australian Research Council Future Fellowship grant FT140100865. This research was undertaken with the assistance of resources and services from the Queensland Cyber Infrastructure Foundation (QCIF) and the National Computational Infrastructure (NCI), which is supported by the Australian Government.

## Word Counts

This section is *not* included in the word count.

### Notes on Nature Methods Brief Communication

Abstract: 3 sentences, 70 words.

Main text: 3 pages, 2 figures, 1000-1500 words, more figures possible if under 3 pages

### Statistics on word count

~~~
File: Article.tex
Encoding: utf8
Sum count: 8386
Words in text: 7889
Words in headers: 61
Words outside text (captions, etc.): 426
Number of headers: 21
Number of floats/tables/figures: 7
Number of math inlines: 9
Number of math displayed: 1
Subcounts:
text+headers+captions (#headers/#floats/#inlines/#displayed)
61+13+0 (1/0/0/0) _top_
249+1+0 (1/0/0/0) Abstract
11+0+0 (0/0/0/0) main
1025+1+66 (1/1/0/0) Section: Introduction
0+1+0 (1/0/0/0) Section: Methods
268+2+106 (1/1/0/0) Subsection: Network Architecture
111+2+0 (1/0/3/1) Subsection: Dipole Kernel
170+2+0 (1/0/2/0) Subsection: Training Procedure
467+9+63 (5/1/2/0) Subsection: Test
75+1+0 (1/0/0/0) Section: Results
74+4+34 (1/1/0/0) Subsection: Experiment 1: Synthetic Data
78+4+46 (1/1/0/0) Subsection: Experiment 2: Forward Simulation
118+9+66 (1/1/0/0) Subsection: Experiment 3: Single Orientation 3D GRE MRI Phase Data
46+8+45 (1/1/0/0) Subsection: Experiment 4: Clinical 3D GRE MRI Phase Data
861+1+0 (1/0/1/0) Section: Discussion
215+1+0 (1/0/0/0) Section: Conclusions
4060+2+0 (2/0/1/0) Section: Bibliography
~~~

